# Improved accuracy for automated counting of a fish in baited underwater videos for stock assessment

**DOI:** 10.1101/2021.02.01.429285

**Authors:** RM Connolly, DV Fairclough, EL Jinks, EM Ditria, G Jackson, S Lopez-Marcano, AD Olds, KI Jinks

## Abstract

The ongoing need to sustainably manage fishery resources necessitates fishery-independent monitoring of the status of fish stocks. Camera systems, particularly baited remote underwater video stations (BRUVS), are a widely-used and repeatable method for monitoring relative abundance, required for building stock assessment models. The potential for BRUVS-based monitoring is restricted, however, by the substantial costs of manual data extraction from videos. Computer vision, in particular deep learning models, are increasingly being used to automatically detect and count fish at low abundances in videos. One of the advantages of BRUVS is that bait attractants help to reliably detect species in relatively short deployments (e.g. 1 hr). The high abundances of fish attracted to BRUVS, however, make computer vision more difficult, because fish often occlude other fish. We build upon existing deep learning methods for identifying and counting a target fisheries species across a wide range of fish abundances. Using BRUVS imagery targeting a recovering fishery species, Australian snapper (*Chrysophrys auratus*), we tested combinations of three further mathematical steps likely to generate accurate, efficient automation: 1) varying confidence thresholds (CTs), 2) on/off use of sequential non-maximum suppression (Seq-NMS), and 3) statistical correction equations. Output from the deep learning model was accurate at very low abundances of snapper; at higher abundances, however, the model over-predicted counts by as much as 50%. The procedure providing the most accurate counts across all fish abundances, with counts either correct or within 1 to 2 of manual counts (R^2^ = 93.4%), used Seq-NMS, a 55% confidence threshold, and a cubic polynomial corrective equation. The optimised modelling provides an automated procedure offering an effective and efficient method for accurately identifying and counting snapper in BRUV footage. Further testing is required to ensure that automated counts of snapper remain accurate in the survey region over time, and to determine the applicability to other regions within the distributional range of this species. For monitoring stocks of fishery species more generally, the specific equations will differ but the procedure demonstrated here would help to increase the usefulness of BRUVS, while decreasing costs.

## 1 Introduction

The ongoing need to maximise fishery harvests while maintaining stocks at sustainable levels demands efficient in-water monitoring of fish abundances. The advent of robust yet inexpensive underwater cameras provides a potential step-change in increased efficiency of monitoring stocks. Unfortunately, the requirement for manual processing of underwater videos to count target species severely curtails the scalability of camera systems (Sheaves et al. 2020). Automated image analysis can overcome this bottleneck, but technical limitations have restricted its use for routine fisheries monitoring to date (Lopez-Marcano et al. 2021, Tseng & Kuo 2020, Yang et al. 2020).

The most common measure of fish abundance derived from underwater videos is the maximum count of a target species in any one frame (MaxN). A single MaxN per video is the most commonly reported metric (Whitmarsh et al. 2016), but multiple MaxN measures over short time intervals of video, and averages of these, are recommended as being more reliable statistically (Schobernd et al. 2014). By removing the costly step of manual counting, automation can encourage extraction of more values for video and thus greater statistical rigour. Automated analysis needs to extract these values accurately and efficiently to be useful.

Baited remote underwater video stations (BRUVS) are the most widely used application of videos for monitoring fish abundances (Whitmarsh et al. 2016), and automated analysis therefore needs to be accurate specifically for this method. Along with issues common to all underwater image analysis, such as variable water clarity and complex, dynamic backgrounds (Siddiqui et al. 2019, Yang et al. 2020), the BRUVS technique raises another challenge by generating potentially large ranges of fish abundances, from none to many individual fish. Automated analysis needs to report accurately across this wide range of abundances, overcoming significant occlusion issues (where an individual fish can obscure parts of another fish) at higher fish densities.

Efficient, automated identification and counting of fish in underwater images has become possible with the development of computer vision, in particular deep learning (DL), a form of machine learning that automatically extracts features from raw imagery (Le Cun et al. 2015). DL has been used for classification of individual fish images into species classes (Salman et al. 2016, Siddiqui et al. 2018, dos Santos & Goncalves 2019), and for object detection *and* classification on underwater video streams (Mandal et al. 2018, Villon et al. 2018). In unbaited remote underwater video stations, video analysis has been successful on individual target species (Ditria et al. 2020 a) and multiple species selected from fish assemblages (Villon et al. 2018, Villon et al. 2020, Knausgard et al. 2020). Here, we build upon existing DL methods for fish identification and counting to improve accuracy for BRUVS over a wide range of fish abundances. The objective is to automate the current manual analysis of BRUVS videos for monitoring abundances of a key fisheries species, *Chrysophrys auratus* (Australasian snapper, family Sparidae). This popular recreational and commercial species has suffered stock declines and corresponding management responses across much of the species distribution in Australia (Fowler et al. 2018). In Western Australia, there is renewed focus on the development of fishery-independent methods for monitoring relative abundance over time as an input to stock assessment models.

## 2 Materials and Methods

To achieve our aim of building on output from existing DL models to improve counts of target species in BRUVS, we first trained a DL model on footage from BRUVS deployed to monitor abundances of Australasian snapper (snapper) off Shark Bay in the Gascoyne region of Western Australia. We then applied combinations of mathematical procedures to improve accuracy of automated counts.

### Dataset and deep learning model

The stock of snapper in oceanic waters off Shark Bay (∼26°S) in the Gascoyne region of Western Australia was recently assessed (2017) as being at high risk with a range of management actions subsequently introduced in 2018 to reduce exploitation and assist stock recovery (Jackson et al. 2020). Fishery-independent monitoring of snapper at breeding aggregation sites off Shark Bay is in its infancy, with underwater camera systems being tested for future surveys. We used video footage from the six monitoring sites surveyed following the introduction of fishery spatial closures. BRUVS used the standard set-up described by Langlois et al. (2020), and were deployed on mixed rock-sand habitats in 30-60 m water depth, set for 1 hr at each site.

Each video was divided into two segments, one used for training, and one for testing. For training, footage from 10 – 60 minutes into each video was used. We selected numerous snips within this period to provide a wide range of snapper densities and presence of other species. Individual snapper were identified manually by one of the authors (KIJ), who is an experienced fish biologist, and manually annotated with bounding boxes (following Ditria et al. 2020 b). The annotator could play videos back and forth to obtain different views of individual fish to increase confidence in snapper detections. In all, 4,300 individual snapper were annotated. Importantly, this included fish at all angles to the camera, and parts of fish (e.g. head only) where the remainder of the individual was unobservable, either obscured by other fish or outside the field of view. In early trials, we noted that two other species superficially resembling snapper (brown-stripe snapper, *Lutjanus vitta*, and stripey snapper, *Lutjanus carponotatus*) occasionally caused misidentification (i.e. false positive labelling as *Chrysophrys auratus*), and we therefore also annotated and trained the models for these species (50 and 135 annotations, respectively).

For testing, we used the first 5 minutes of videos from each site. This selection presented an appropriately wide range of fish densities, since BRUVS typically have no fish present as the camera drop begins, with high densities by the 5 min mark as the bait attractant takes effect (Whitmarsh et al. 2016). In these test video segments, the number of snapper (N) was recorded every 30 seconds (i.e. 10 records over 5 minutes) by the same observer as for training. These manually extracted N values provided ground-truth results against which computer predictions were tested.

We used a convolutional neural network framework for object detection, specifically an implementation of Faster R-CNN developed by Massa and Girshick (2018). Model development was conducted using a ResNet50 configuration, pre-trained on the ImageNet-1k dataset. This method successfully detects and counts target species at low densities in unbaited RUVs (Ditria et al. 2020 a). Model training, prediction, and testing tasks were conducted on a Microsoft Azure Data Science Virtual Machine powered by an NVIDIA V100 GPU. Overfitting was minimised using the early-stopping technique (Prechelt, 1998).

### Mathematical procedures

In seeking to improve the accuracy of DL model output, we applied mathematical procedures to raw computer predictions. We tested numerous combinations of three key mathematical components considered likely to generate accurate, efficient automation: 1) varying confidence thresholds (CTs), 2) on/off use of sequential non-maximum suppression (Seq-NMS), and 3) statistical correction equations. Seq-NMS was tried both before and after varying CTs. Statistical correction equations were always applied last. We also tried variations of other aspects, such as image resolution, but these did not provide measurable improvement and are not reported on further. Selection of CTs, the values above which objects are classified into a class (here, snapper), are an important determinant in balancing false positive and false negatives (and, therefore, in maximising true positives). We tried CTs from 0 – 95% in 5% increments. Seq-NMS is a spatio-temporal filter that creates detection chains by analysing neighbouring frames (Han et al. 2016). It is regarded as a useful procedure where DL models are over-predicting, which in initial trials on snapper BRUVS we identified as an issue at high fish densities. As a final mathematical component, we applied corrective equations to output from combinations of CT and Seq-NMS. Given the patterns of errors in predictions we most commonly observed, we tried linear, quadratic and cubic polynomial equations with randomly varying constants. In total, 120 combinations of the three components were tested (combinations of 20 CTs, Seq-NMS on/off, 3 forms of equations). In all cases, the measure of accuracy was the fit of computer predictions of N against ground-truth values, across the entire range of fish densities (quantified by R^2^ value).

## 3 Results

Raw automated predictions were generally inaccurate, particularly at high snapper abundances (Fig. 1). Automated counts were close to ground-truth counts only when actual fish counts were very low (e.g. 0 - 3 fish). At higher abundances, raw predictions were considerably greater than ground-truth values (Fig. 1). This inaccuracy became progressively worse at increasing abundances, so that at maximal ground-truth values of approximately 30 fish, raw predictions were on average about 50% above those (approx. 45 fish). The over-prediction was almost solely due to high numbers of false positive detections of snapper, predominantly as double and triple detection of the same individual fish; for example, the head and tail of one fish were counted as two fish (see example in Supplementary material, Fig. S1)

**Figure 1.**
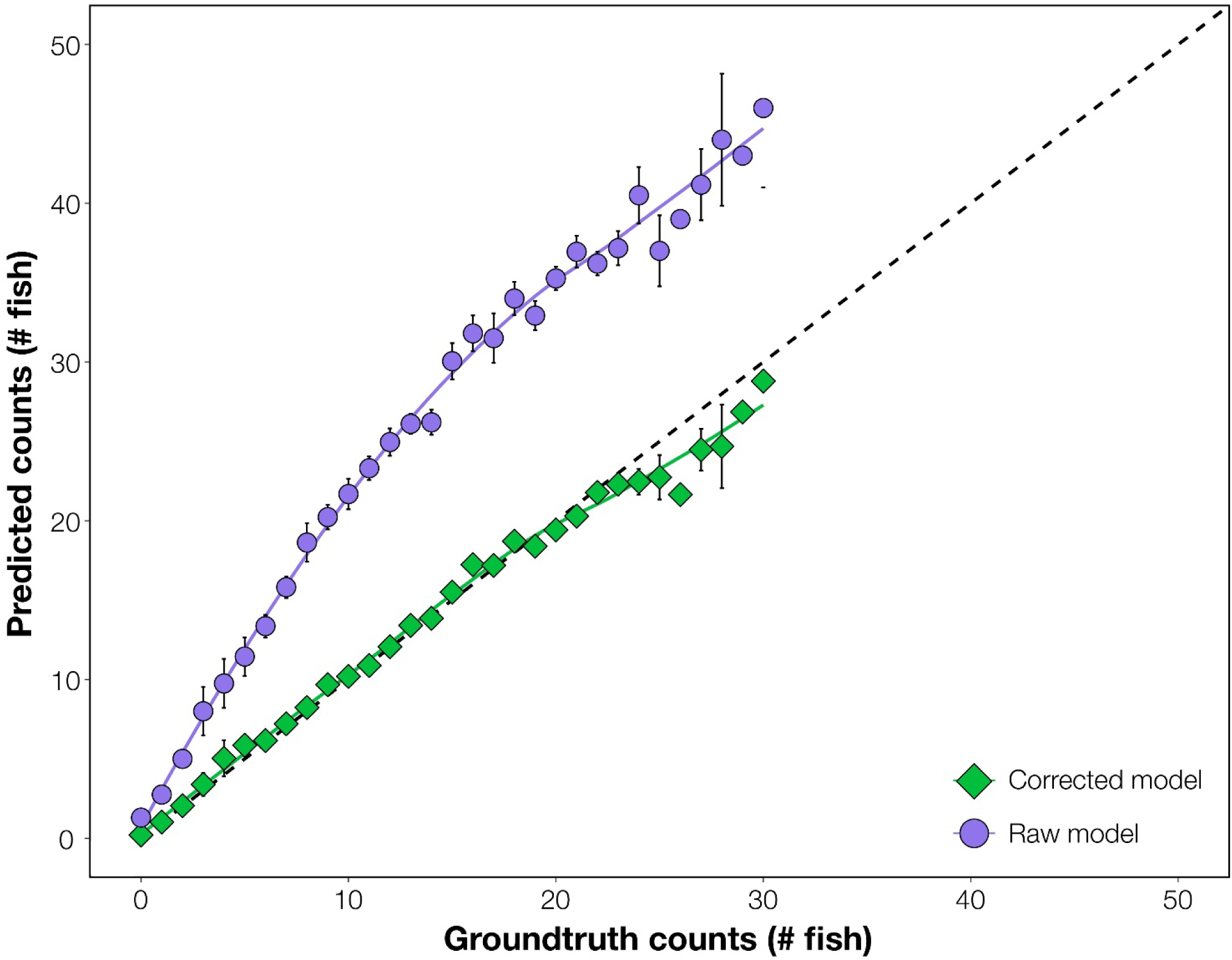
Comparison of accuracy of automated counts of the target species, Australasian snapper, from the raw deep learning output (Raw model) and the enhanced modelling procedure (Corrected model). In both cases, prediction counts are tested for accuracy against ground-truth (manual) counts, with more accurate results closer to the 1:1 dashed line. Raw model counts are unmodified from the deep learning model. Corrected model counts use automated mathematical enhancements – Seq-NMS, optimum confidence threshold, and corrective equation - on the deep learning output. Data are combined from all sites. Error bars = SE.

After optimising combinations of mathematical procedures, computer predictions became highly accurate (Fig. 1). Corrected predictions were on average the same as ground-truth values at all fish abundances, with only slight under-prediction at ground-truthed abundances above 25 fish (under-predicted by 1 – 2 fish). The optimal enhancements were, in order: 1) Seq-NMS on, 2) CT of 55%, and 3) a cubic polynomial corrective equation applied (N’ = A + B.N + C.N^2^ + D.N^3^ where A = 12.9, B = 132.6, C = 14.4, D = −3.0). This optimum was selected at the highest R^2^ from the 120 combinations (Table S1). A comparison of automated counts from the final revised modelling procedure against manual ground-truth counts gave an R^2^ of 93.4%.

The effectiveness of the optimum model procedure is illustrated in predictions at each 30 seconds over the first five minutes of BRUVS drops, shown for sites selected for their strikingly different snapper abundances (Fig. 2). The MaxN values and the average MaxN values from computer generated predictions were very similar to ground-truth values at each time interval, i.e. either exactly the same or within 1 or 2 of actual counts (Fig. 2).

**Figure 2.**
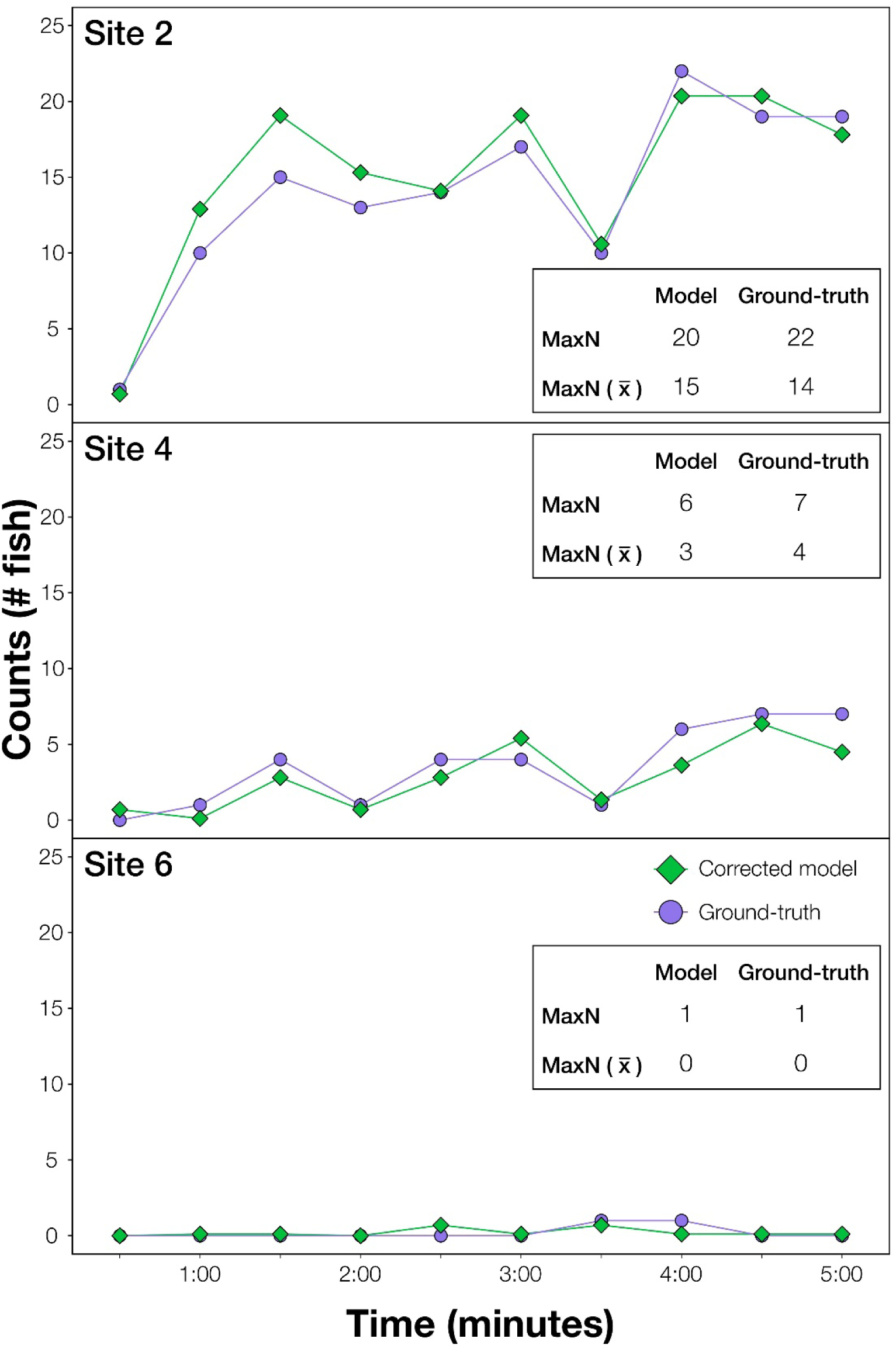
Results illustrating the effectiveness of the modelling procedure at automatically extracting counts of the target species, Australasian snapper, from videos. Sites shown are a selection demonstrating accuracy at high (Site 2), medium (Site 4) and very low (Site 6) snapper abundances. At each site, ground-truth and modelled (optimised model using deep learning and additional operations) counts are shown every 30 seconds for the first 5 minutes of video. This imagery was independent of that used in training. MaxN is the greatest fish count over the 5 min period, and MaxN 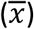 is the average MaxN over the 5 min period.

## 4 Discussion

The refined procedure of deep learning with additional automated mathematical operations produced a highly effective and efficient method for identifying and counting target fish in BRUVS. The processing procedure provides rapid, automated extraction of snapper counts from zero to high abundances. The final optimised procedure utilised a combination of Seq-NMS, a specific confidence threshold, and a cubic polynomial corrective equation (Fig. 3). The automated processing procedure will be useful for snapper monitoring, and will encourage expansion of monitoring sites and times while avoiding increased costs of manual processing. It will also encourage reporting of MaxN values at much more frequent intervals within BRUVS videos, an important aspect of increasing the rigour of fisheries monitoring (Schobernd et al. 2014).

**Figure 3.**
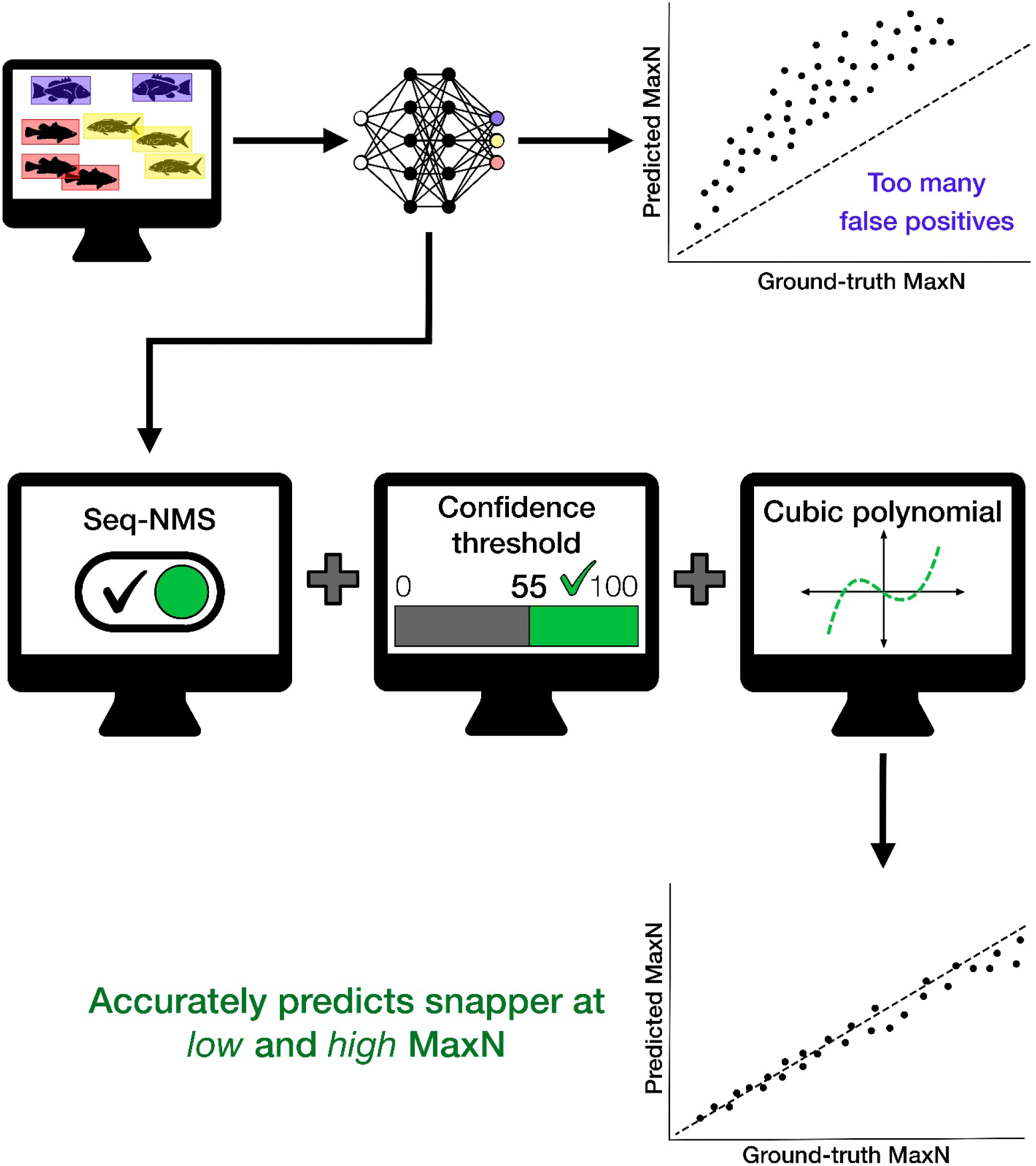
Conceptual diagram summarising steps used to create the deep learning model and apply additional mathematical operations to improve accuracy of automated counts of the target species, Australasian snapper, particularly at higher fish densities. The optimal procedure uses Seq-NMS, a confidence threshold of 55%, and a cubic polynomial corrective equation, and produces the accurate automated counts shown in Fig. 1.

The procedure demonstrated here can stimulate improvements in automation of BRUVS data extraction more generally, and be used for BRUVS automation for many different species and situations. For species other than snapper, selected CTs and the form of corrective equation can be expected to vary. Even within a single dataset, the optimum CT is known to differ among species and with different amounts of training data (Villon et al. 2020).

Although the demonstrated procedure has been extremely successful, we suggest some caveats and recommendations for further trials and testing. Two common challenges in computer vision, domain shift and concept drift, will need to be addressed when applying our model more generally. Our model performance depends on the environment, or domain, in which it was trained (Ditria et al. 2020 b) At this stage, the optimal procedure is addressed only for snapper at the multiple sites within the survey area in the Gascoyne region of Western Australia. The usefulness of the model will ultimately need demonstrating in different regions across the distributional range of snapper. The model will need checking in other places where aspects such as habitat backgrounds and species assemblages will vary - a concept known as domain shift (Kalogeiton et al. 2016). It will also require testing in situations where the size composition of snapper differs from that in this study. For example, BRUVs are used in monitoring the abundance of snapper recruits on the lower west coast of Australia (Wakefield et al., 2013). We expect that use of the model in other places will require different CTs or corrective equations, although it is also possible that a more generic model might be developed that is effective across the snapper distributional range. Our model is also linked to a specific time, and so further work is required to guard against inaccuracies due to concept drift (Hashmani et al. 2019). Changes in the environment or camera equipment over time can reduce the accuracy of computer vision models (e.g. Langenkamper et al. 2020). The current study produced a very accurate procedure for videos from the initial survey of the Gascoyne region. It will need testing over time as videos become available from future monitoring surveys, to address aspects of the environment likely to change, such as known tropicalisation of reefal habitats in Western Australia (Arias-Ortiz et al. 2018).

There is clear potential for DL automation to revolutionise in-water monitoring of animal abundances (Christin et al. 2019). The applications of computer vision to fisheries science are at the early stages of being realised (Lopez-Marcano et al. 2021). BRUVS are already used for safe and repeatable monitoring of fish abundances in a range of situations (Harvey et al. in press), and we hope that the procedures demonstrated here can increase the usefulness of BRUVS, while decreasing costs, and ultimately improving fishery-independent stock assessments.

## 5 Data Availability Statement

The dataset generated for this study is available on request to the corresponding author. Code for the mathematical enhancements will be made available on Github at the time of publication.

## 6 Acknowledgments

The authors thank technical staff (particularly Nick Jarvis) of the Department of Primary Industries and Regional Development for conducting video sampling and skippers and owner of the FV Ada Clara. The study benefited from collaborations with all researchers in the Global Wetlands Project.

---

**Figure S1.**
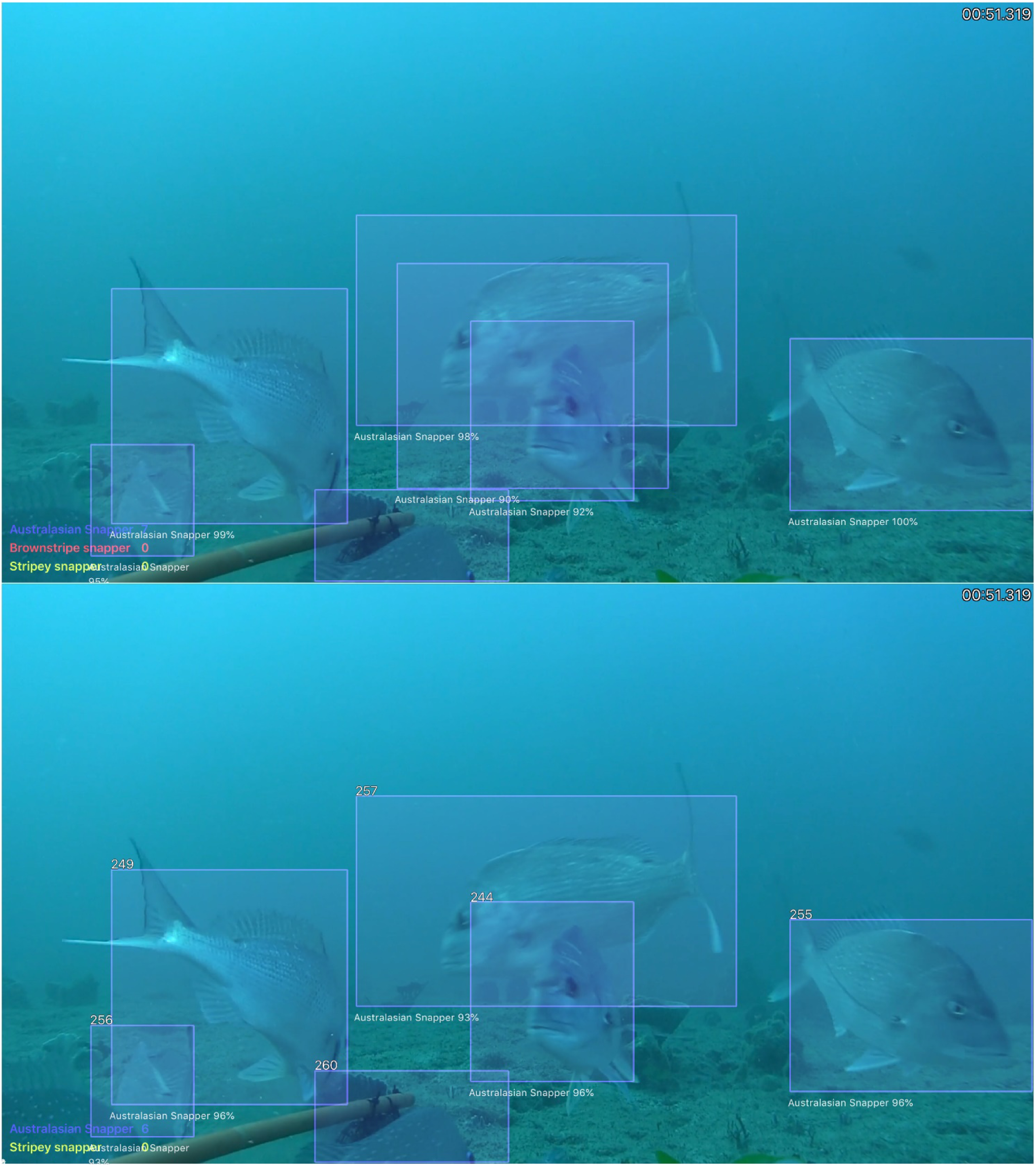
Examples of computer detections of Australasian snapper. Top panel shows inaccurate predictions, specifically numerous false positive detections. Bottom panel shows accurate detections.

**Table S1.** Closeness of fit of predicted counts against ground-truth counts measured as R^2^ compared with 1:1 line. Polynomial level: 1 = linear, 2 = square, 3 = cubic.

| **Confidence**<br>**Threshold**<br>**(%)** | **Seq-NMS**<br>**Y = on**<br>**N = off** | **Polynomial**<br>**1,2,3** | **Fit**<br>**($R^2$)** |
| --- | --- | --- | --- |
| 0 | N | 1 | 0.870 |
| 0 | N | 2 | 0.871 |
| 0 | N | 3 | 0.874 |
| 0 | Y | 1 | 0.917 |
| 0 | Y | 2 | 0.927 |
| 0 | Y | 3 | 0.927 |
| 5 | N | 1 | 0.87 |
| 5 | N | 2 | 0.871 |
| 5 | N | 3 | 0.874 |
| 5 | Y | 1 | 0.917 |
| 5 | Y | 2 | 0.927 |
| 5 | Y | 3 | 0.927 |
| 10 | N | 1 | 0.870 |
| 10 | N | 2 | 0.871 |
| 10 | N | 3 | 0.874 |
| 10 | Y | 1 | 0.917 |
| 10 | Y | 2 | 0.927 |
| 10 | Y | 3 | 0.927 |
| 15 | N | 1 | 0.871 |
| 15 | N | 2 | 0.872 |
| 15 | N | 3 | 0.875 |
| 15 | Y | 1 | 0.915 |
| 15 | Y | 2 | 0.925 |
| 15 | Y | 3 | 0.925 |
| 20 | N | 1 | 0.880 |
| 20 | N | 2 | 0.882 |
| 20 | N | 3 | 0.883 |
| 20 | Y | 1 | 0.918 |
| 20 | Y | 2 | 0.927 |
| 20 | Y | 3 | 0.928 |
| 25 | N | 1 | 0.886 |
| 25 | N | 2 | 0.889 |
| 25 | N | 3 | 0.890 |
| 25 | Y | 1 | 0.918 |
| 25 | Y | 2 | 0.927 |
| 25 | Y | 3 | 0.927 |
| 30 | N | 1 | 0.889 |
| 30 | N | 2 | 0.892 |
| 30 | N | 3 | 0.893 |
| 30 | Y | 1 | 0.917 |
| 30 | Y | 2 | 0.927 |
| 30 | Y | 3 | 0.927 |
| 35 | N | 1 | 0.898 |
| 35 | N | 2 | 0.903 |
| 35 | N | 3 | 0.904 |
| 35 | Y | 1 | 0.917 |
| 35 | Y | 2 | 0.926 |
| 35 | Y | 3 | 0.926 |
| 40 | N | 1 | 0.903 |
| 40 | N | 2 | 0.909 |
| 40 | N | 3 | 0.909 |
| 40 | Y | 1 | 0.918 |
| 40 | Y | 2 | 0.927 |
| 40 | Y | 3 | 0.927 |
| 45 | N | 1 | 0.904 |
| 45 | N | 2 | 0.909 |
| 45 | N | 3 | 0.910 |
| 45 | Y | 1 | 0.923 |
| 45 | Y | 2 | 0.932 |
| 45 | Y | 3 | 0.932 |
| 50 | N | 1 | 0.907 |
| 50 | N | 2 | 0.912 |
| 50 | N | 3 | 0.913 |
| 50 | Y | 1 | 0.923 |
| 50 | Y | 2 | 0.932 |
| 50 | Y | 3 | 0.932 |
| 55 | N | 1 | 0.905 |
| 55 | N | 2 | 0.911 |
| 55 | N | 3 | 0.912 |
| 55 | Y | 1 | 0.923 |
| 55 | Y | 2 | 0.934 |
| 55 | Y | 3 | 0.934 |
| 60 | N | 1 | 0.904 |
| 60 | N | 2 | 0.910 |
| 60 | N | 3 | 0.912 |
| 60 | Y | 1 | 0.921 |
| 60 | Y | 2 | 0.932 |
| 60 | Y | 3 | 0.933 |
| 65 | N | 1 | 0.905 |
| 65 | N | 2 | 0.911 |
| 65 | N | 3 | 0.914 |
| 65 | Y | 1 | 0.920 |
| 65 | Y | 2 | 0.931 |
| 65 | Y | 3 | 0.931 |
| 70 | N | 1 | 0.910 |
| 70 | N | 2 | 0.920 |
| 70 | N | 3 | 0.920 |
| 70 | Y | 1 | 0.912 |
| 70 | Y | 2 | 0.921 |
| 70 | Y | 3 | 0.921 |
| 75 | N | 1 | 0.907 |
| 75 | N | 2 | 0.916 |
| 75 | N | 3 | 0.917 |
| 75 | Y | 1 | 0.904 |
| 75 | Y | 2 | 0.911 |
| 75 | Y | 3 | 0.912 |
| 80 | N | 1 | 0.907 |
| 80 | N | 2 | 0.914 |
| 80 | N | 3 | 0.917 |
| 80 | Y | 1 | 0.887 |
| 80 | Y | 2 | 0.895 |
| 80 | Y | 3 | 0.896 |
| 85 | N | 1 | 0.901 |
| 85 | N | 2 | 0.907 |
| 85 | N | 3 | 0.912 |
| 85 | Y | 1 | 0.852 |
| 85 | Y | 2 | 0.853 |
| 85 | Y | 3 | 0.858 |
| 90 | N | 1 | 0.883 |
| 90 | N | 2 | 0.890 |
| 90 | N | 3 | 0.896 |
| 90 | Y | 1 | 0.612 |
| 90 | Y | 2 | 0.623 |
| 90 | Y | 3 | 0.624 |
| 95 | N | 1 | 0.830 |
| 95 | N | 2 | 0.832 |
| 95 | N | 3 | 0.848 |
| 95 | Y | 1 | 0.183 |
| 95 | Y | 2 | 0.250 |
| 95 | Y | 3 | 0.292 |

## References

Arias-Ortiz, A., Serrano, O., Masque, P., Lavery, P.S., Mueller, U., et al. (2018). A marine heatwave drives massive losses from the world’s largest seagrass carbon stocks. Nat. Clim. Change. 8, 338–344.

Christin, S., Hervet, É., and Lecomte, N. (2019). Applications for deep learning in ecology. Methods Ecol. Evol. 10, 1632–1644.

Ditria, E.M., Sievers, M., Connolly, R.M., Jinks, E.L., and Lopez-Marcano, S. (2020). Deep learning for automated analysis of fish abundance: the benefits of training across multiple habitats. Environ. Monit. Assess. 192, 698.

Ditria, E.M., Lopez-Marcano, S., Sievers, M., Jinks, E.L., Brown, C.J., and Connolly, R.M. (2020). Automating the analysis of fish abundance using object detection: Optimizing animal ecology with deep learning. Front. Mar. Sci. 7, 429.

dos Santos, A.A., and Gonçalves, W.N. (2019). Improving Pantanal fish species recognition through taxonomic ranks in convolutional neural networks. Ecol. Inform. 53, 100977.

Fowler, A., Jackson, G., Stewart, J., Hamer, P., and Roelofs, A. (2018). Snapper, Chrysophrys auratus, in status of key Australian fish stocks reports 2018. Fisheries Research and Development Corporation (FRDC). Canberra.

Han, W., Khorrami, P., Paine, T.L., Ramachandran, P., Babaeizadeh, M., Shi, H., et al. (2016). Seq-NMS for video object detection. International Conference on Learning Representations. (ICLR). Peurto Rico, USA.

Harvey, E.S., Mclean, D.L., Goetze, J.S., Saunders, B.J., Langlois, T.J., Monk, J., et al. (in press). The BRUVS workshop - an Australian-wide synthesis of baited remote underwater video data to answer broad-scale ecological questions about fish, sharks and rays. Mar. Policy.

Hashmani, M.A., Jameel, S.M., Al-Hussain, H., Rehman, M., and Budiman, A. (2019). Accuracy performance degradation in image classification models due to concept drift. Int. J. Adv. Comput. Sci. Appl. (IJACSA). 10(5).

Jackson, G., Denham, A., Hesp, A., Hall, N., Fisher, E., and Stephenson, P. (2020). Gascoyne demersal scalefish resource. Department of Primary Industries and Regional Development (DPIRD) report 009. Perth, Western Australia.

Kalogeiton, V., Ferrari, V., and Schmid, C. (2016). Analysing domain shift factors between videos and images for object detection. IEEE Trans. Pattern Anal. Mach. Intell. 38, 2327–2334.

Knausgård, K.M., Wiklund, A., Sørdalen, T.K., Halvorsen, K., Kleiven, A.R., Jiao, L., et al. (2020). Temperate fish detection and classification: A deep learning based approach. arXiv:2005.07518

Langenkämper, D., van Kevelaer, R., Purser, A., and Nattkemper, T.W. (2020). Gear-induced concept drift in marine images and its effect on deep learning classification. Front. Mar. Sci. 7, 506.

Langlois, T., Goetze, J., Bond, T., Monk, J., Abesamis, R.A., Asher, J., et al. (2020). A field and video annotation guide for baited remote underwater stereo-video surveys of demersal fish assemblages. Methods Ecol. Evol. 11, 1401–1409.

LeCun, Y., Bengio, Y., and Hinton, G. (2015). Deep learning. Nature. 521, 436–444.

Lopez-Marcano, S., Brown, C.J., Sievers, M., and Connolly, R.M. (2021). The slow rise of technology: Computer vision techniques in fish population connectivity. Aquat. Conserv.: Mar. Freshwat. Ecosyst. 31, 210–217.

Mandal, R., Connolly, R.M., Schlacher, T.A., and Stantic, B. (2018). Assessing fish abundance from underwater video using deep neural networks, 2018 International Joint Conference on Neural Networks (IJCNN), 1–6.

Massa, F., and Girshick, R. (2018). maskrcnn-benchmark: Fast, modular reference implementation of Instance Segmentation and Object Detection algorithms in PyTorch. https://github.com/facebookresearch/maskrcnn-benchmark, 2018.

Prechelt, L. (1998). “Early stopping-but when?,” in Neural Networks: Tricks of the trade. (Berlin Heidelberg: Springer), 55–69.

Salman, A., Jalal, A., Shafait, F., Mian, A., Shortis, M., Seager, J., et al. (2016). Fish species classification in unconstrained underwater environments based on deep learning. Limnol. Oceanogr. Methods. 14, 570–585.

Schobernd, Z.H., Bacheler, N.M., and Conn, P.B. (2014). Examining the utility of alternative video monitoring metrics for indexing reef fish abundance. Can. J. Fish. Aquat Sci. 71, 464–471.

Sheaves, M., Bradley, M., Herrera, C., Mattone, C., Lennard, C., Sheaves, J., et al. (2020). Optimizing video sampling for juvenile fish surveys: Using deep learning and evaluation of assumptions to produce critical fisheries parameters. Fish Fish. 21, 1259–1276.

Siddiqui, S.A., Salman, A., Malik, M.I., Shafait, F., Mian, A., Shortis, M.R., et al. (2018). Automatic fish species classification in underwater videos: exploiting pre-trained deep neural network models to compensate for limited labelled data. ICES J. Mar. Sci. 75, 374–389.

Tseng, C.-H., and Kuo, Y.-F. (2020). Detecting and counting harvested fish and identifying fish types in electronic monitoring system videos using deep convolutional neural networks. ICES J. Mar. Sci. 77, 1367–1378.

Villon, S., Mouillot, D., Chaumont, M., Darling, E.S., Subsol, G., Claverie, T., et al. (2018). A deep learning method for accurate and fast identification of coral reef fishes in underwater images. Ecol. Inform. 48, 238–244.

Villon, S., Mouillot, D., Chaumont, M., Subsol, G., Claverie, T., and Villéger, S. (2020). A new method to control error rates in automated species identification with deep learning algorithms. Sci. Rep. 10, 1–13.

Wakefield, C.B., Lewis, P.D., Coutts, T.B., Fairclough, D.V., and Langlois, T.J. (2013). Fish assemblages associated with natural and anthropogenically-modified habitats in a marine embayment: Comparison of baited videos and opera-house traps. PLoS. ONE. 8, e59959.

Whitmarsh, S.K., Fairweather, P.G., and Huveneers, C. (2017). What is Big BRUVver up to? Methods and uses of baited underwater video. Rev. Fish Biol. Fish. 27, 53–73.

Yang, L., Liu, Y., Yu, H., Fang, X., Song, L., Li, D., et al. (2020). Computer Vision Models in Intelligent Aquaculture with Emphasis on Fish Detection and Behavior Analysis: A Review. Arch. Comput. Methods Eng. 1–32.

